# The role of *a priori*-identified addiction and smoking gene sets in smoking behaviors

**DOI:** 10.1101/732321

**Authors:** Luke M. Evans, Emma C. Johnson, Whitney E. Melroy-Grief, John K. Hewitt, Charles A. Hoeffer, Matthew C. Keller, Laura M. Saba, Jerry A. Stitzel, Marissa A. Ehringer

## Abstract

**Introduction:** Smoking is a leading cause of death, and genetic variation contributes to smoking behaviors. Identifying genes and sets of genes that contribute to risk for addiction is necessary to prioritize targets for functional characterization and for personalized medicine.

**Methods:** We performed a gene set-based association and heritable enrichment study of two addiction-related gene sets, those on the Smokescreen Genotyping Array and the nicotinic acetylcholine receptors, using the largest available GWAS summary statistics. We assessed smoking initiation, cigarettes per day, smoking cessation, and age of smoking initiation.

**Results:** Individual genes within each gene set were significantly associated with smoking behaviors. Both sets of genes were significantly associated with cigarettes per day, smoking initiation, and smoking cessation. Age of initiation was only associated with the Smokescreen gene set. While both sets of genes were enriched for trait heritability, each accounts for only a small proportion of the SNP-based heritability (2-12%).

**Conclusions:** These two gene sets are associated with smoking behaviors, but collectively account for a limited amount of the genetic and phenotypic variation of these complex traits, consistent with high polygenicity.

**Implications:** We evaluated evidence for association and heritable contribution of expert-curated and bioinformatically identified sets of genes related to smoking. Although they impact smoking behaviors, these specifically targeted genes do not account for much of the heritability in smoking and will be of limited use for predictive purposes. Advanced genome-wide approaches and integration of other ‘omics data will be needed to fully account for the genetic variation in smoking phenotypes.

## INTRODUCTION

Smoking is one of the most prominent causes of death in the United States, leading to numerous diseases and shortened life expectancy^1^. While the majority of smokers report a desire to quit, very few are able^2^. Furthermore, although smoking rates have decreased, other forms of nicotine consumption are rapidly growing, such as adolescent vaping^3^, demonstrating a pressing need to characterize the underlying biology of nicotine use and smoking to reduce subsequent premature death.

Abundant evidence indicates that up to 50% of the phenotypic variance in nicotine dependence is due to genetic factors^4–6^. In addition to this evidence from twin, adoption and family studies, genome-wide association studies (GWAS) have begun to identify variants associated with smoking behaviors^7,8^, providing insights into the genetic etiology of smoking and nicotine dependence. A key finding from such studies has been the high polygenicity of these traits – Liu et al.^7^ found over two hundred conditionally independent loci throughout the genome that influenced smoking initiation, escalation, and cessation using over 1.2 million individuals, with additional loci expected to be identified as sample sizes increase.

Efficient genotyping and disease-specific arrays have been developed with the aim of identifying particular variants to use in individualized therapies through predictive genetic models^9^. To this end, the Smokescreen Genotyping Array was developed to tag over 1,000 addiction-related genes, identified through expert knowledge, bioinformatic databases, and previous studies^9^. These genes are thought to be strongly associated with addiction, and specifically nicotine use, behaviors.

Nicotinic acetylcholine receptors (nAChRs), which are bound by nicotine, play a key role in smoking behaviors and have been extensively investigated both in human population samples and in functional mouse models. In particular, the nAChR alpha 5 subunit (*CHRNA5*), which is bound by nicotine, is one of the most widely studied genes related to addiction, with replicated GWAS associations^7,8,10^ and functional characterization in the mouse^11,12^. Identifying the key genes that intracellularly interact with them and whether they contribute to heritable variation in smoking phenotypes can lead to biological insight and potentially novel therapeutic targets. Melroy-Greif et al.^13^ performed an extensive literature search, in collaboration with experts in the field of nicotinic acetylcholine receptor (nAChR) research, to identify a set of 107 such genes involved in the upregulation, function, processing, and downstream effects of nAChR signaling (only 36 of these overlap with the Smokescreen Array gene set). Melroy-Greif et al.^13^ posited that this set of genes that are involved in nAChR upregulation, known to occur in reponse to nicotine exposure, play a role in smoking behaviors. While Melroy-Greif et al.^13^ did not find significant gene set associations with smoking phenotypes, GWAS sample sizes have since increased dramatically^7^, leading to greater statistical power to detect such associations.

Understanding which genes and sets of genes are associated with smoking phenotypes may help prioritize future functional studies in model organisms^14^ and so remains an important goal. These two sets of genes provide a starting point to prioritize potential targets, but the high degree of polygenicity of smoking behaviors^7^ raises the question of whether such genes are more strongly associated or enriched than the rest of the genome.

We sought here to assess whether and the degree to which these two sets of genes (“Smokescreen” and “Nicotinic” receptor gene sets) are associated with smoking phenotypes. Using the most recent and largest GWAS summary statistics (GSCAN^7^) for smoking behaviors, we applied gene set association and heritability enrichment analyses to specifically test the contribution of these particular genes to four smoking behaviors: smoking initiation, age of initiation, cigarettes per day, and smoking cessation.

## METHODS

We used Multi-marker Analysis of GenoMic Annotation^15^ (MAGMA) to test gene-level associations of the Smokescreen and Nicotinic gene sets with four smoking behaviors: lifetime smoking initiation, age of smoking initiation, cigarettes per day, and smoking cessation. We used summary statistics from Liu et al.^7^, minus the 23andMe data, and full descriptions of the phenotypes can be found in reference ^7^. We included a control phenotype, alcoholic drinks per week, which was also a GSCAN phenotype. SNP-based genetic correlations^7^ with smoking phenotypes range from 0.1-0.34. While it can be viewed as an addiction-related phenotype^16^, is distinct from smoking behaviors and thus provides an assessment of whether the particular gene sets relate to smoking phenotypes or more generally addiction-related behaviors. These summary statistics correspond to sample sizes of 632,802; 262,990; 263,954; 312,812; and 537,349 individuals for the five phenotypes, respectively.

We applied the competitive MAGMA test to determine whether the gene sets of interest were more strongly associated with these phenotypes than the rest of the genome. We tested two smoking-related gene sets. First, we tested the set of 1,031 addiction genes included on the Smokescreen Genotyping Array^9^ (hereafter referred to as the ‘Smokescreen’ gene set), of which 1,009 were also tagged by variants with GWAS summary statistics from GSCAN. These genes were chosen based on bioinformatic databases and expert curation, being characterized as ‘addiction-relevant’ genes; a full description of the Smokescreen array can be found in Bauerly et al.^9^ The second smoking-relevant gene set was comprised of 107 expert-curated nAChR-related genes that are directly involved in signaling through nAChRs as identified by Melroy-Greif et al.^13^ (hereafter the ‘Nicotinic’ gene set). Thirty-six genes were present in both the Smokescreen and Nicotinic gene sets.

We also tested three control gene sets, that were predicted to show no association with smoking behaviors based on patterns of genetic correlation as presented by Liu et al.^7^: height (444 genes; r_g_ ~ −0.10 – 0.04), Alzheimer’s (476 genes; r_g_ ~ −0.06 – 0.08), and inflammatory bowel disease (234 genes; r_g_ ~ −0.05 – 0.04). These served as negative controls, not expected to have substantial genetic overlap with smoking behaviors. Control sets of genes were identified by querying the GWAS Catalog (https://www.ebi.ac.uk/gwas/) for the terms “height”, “Alzheimer’s”, and “inflammatory bowel disease”, identifying all the gene names tagged by GWAS Catalog as being associated with these traits, and then aggregating these genes into gene sets. As the Nicotinic genes included only protein-coding genes, we excluded pseudogenes, ncRNA genes, lncRNA genes, and miRNA genes from all control gene sets.

In all analyses, variants were annotated to genes using a 25Kb window around the start and end point of each gene in MAGMA in an attempt to include variants that might exert close-range regulatory effects (e.g., within promoter regions) as in previous gene set analyses^17^. We included the default covariates in MAGMA (gene size, density, inverse MAC, per-gene sample size, plus the log value of each) to control for possible confounding factors. We tested five gene sets for each phenotype; therefore, we applied a Bonferroni multiple testing correction for significance for five tests as the within-trait significance threshold (α=0.01).

Following the competitive tests described above, we performed conditional tests of within-trait significant gene sets. These assessed whether the association of a target gene set for a particular phenotype was significant conditional on the effects of additional gene sets. First, we tested the association of the Nicotinic genes conditional on the effects of the Smokescreen genes, which allowed us to determine whether the effects of the Nicotinic genes were simply the effect of genes shared between the two gene sets. Next, we examined the association of the Smokescreen gene set conditional on the Nicotinic gene set. Since no control gene set analysis passed our within-trait Bonferroni significance threshold, we did not evaluate conditional associations of those. In total we performed seven conditional tests and used a Bonferroni correction for seven tests (α=0.00714).

Finally, we performed an enrichment analysis of the heritable contribution of each gene set, relative to the number of markers in the gene set, using partitioned LD score regression^18^ (LDSC). We added annotations for each of the five gene sets listed above to the baseline with LD annotation model^19^, as suggested in the LDSC documentation. We applied a within-trait Bonferroni correction for these enrichment analyses based on five gene sets for each phenotype (α=0.01).

## RESULTS

We found that the smoking gene sets (Smokescreen and Nicotinic) were significantly associated with the four smoking phenotypes, but not the control addiction-related phenotype, drinks per week (Fig. 1, Suppl. Table 1). While both smoking gene sets were significantly associated with cigarettes per day, smoking initiation, and smoking cessation, age of initiation was only associated with the Smokescreen dataset. None of the negative control gene sets was significantly associated with any phenotype (all *p*>0.01).

**Figure 1.**
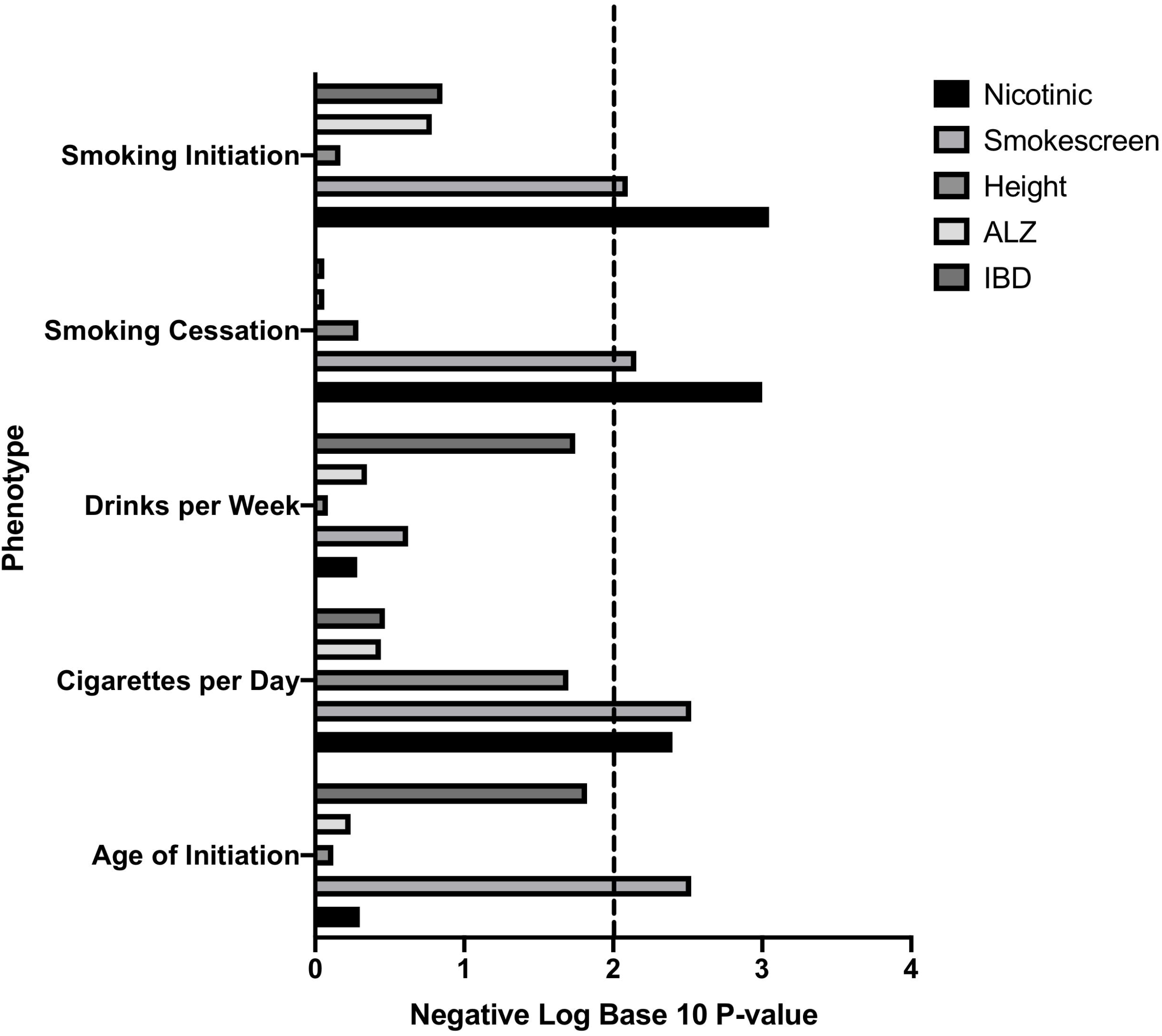
−log10(*p*) of MAGMA competitive tests the four smoking phenotypes and one control trait (Drinks Per Week) across the two smoking gene sets and three control gene sets. Dashed line is thee within-trait Bonferroni correction for multiple testing.

Many genes were present in more than one gene set (36 overlapped between the Nicotinic and Smokescreen gene sets), and 128 individual genes from one or more gene sets were individually significantly associated with one or more phenotypes (Suppl. Table S2). 19 of these were within the Nicotinic gene set, 61 were within the Smokescreen set, and nine of these overlapped (Suppl. Table S2). Thus, though the trait genetic correlations are very weak, some of these genes are likely to influence multiple traits (Suppl. Table S2).

To test whether the smoking gene sets were independently associated with smoking phenotypes, or if the associations for each resulted from the genes overlapping genes in both gene sets, we performed conditional analyses (Table 1). For example, of the 107 Nicotinic genes, 36 overlapped with the Smokescreen gene set. In both cases, the association of Smokescreen and Nicotinic datasets, conditional on the other, remained significant for cigarettes per day (p<0.01). The Nicotinic gene set, conditional on the Smokescreen gene set, was also significantly associated with smoking initiation and cessation, but the Smokescreen gene set, conditional on the Nicotinic gene set, was not (p>0.01).

**Table 1.**
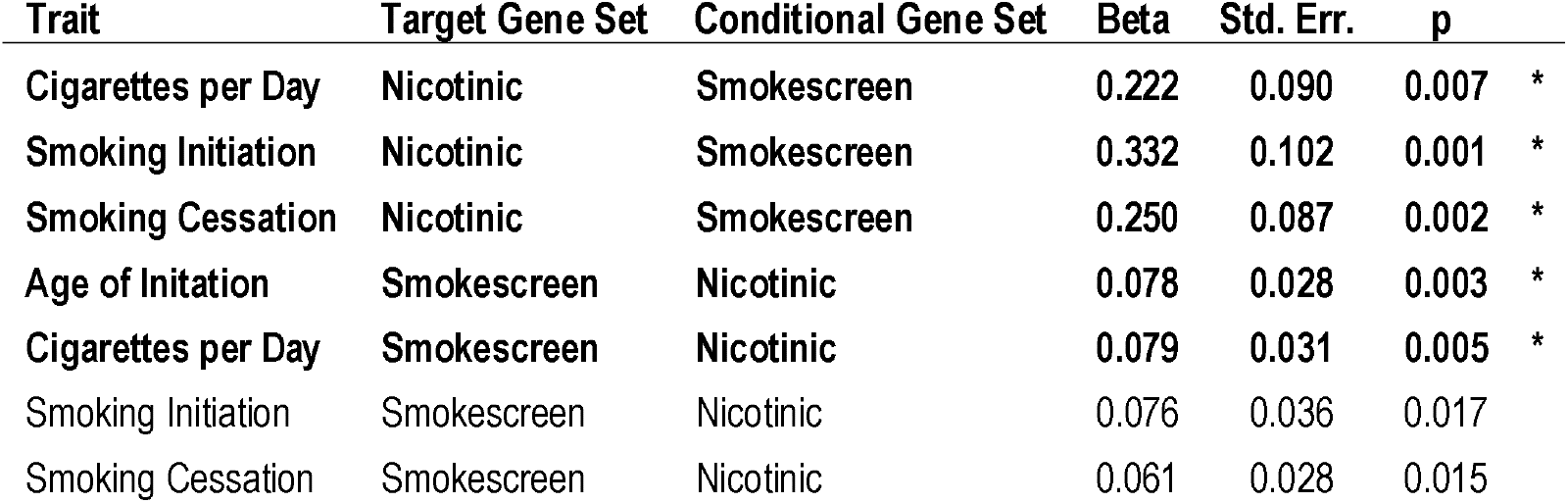
MAGMA conditional test results for the gene sets that were significantly associated with particular traits in the competitive test (Table 1). * indicates p < 0.05/7 < 0.00714.

| Trait | Target Gene Set | Conditional Gene Set | Beta | Std. Err. | p |
| --- | --- | --- | --- | --- | --- |
| Cigarettes per Day | Nicotinic | Smokescreen | 0.222 | 0.090 | 0.007 * |
| Smoking Initiation | Nicotinic | Smokescreen | 0.332 | 0.102 | 0.001 * |
| Smoking Cessation | Nicotinic | Smokescreen | 0.250 | 0.087 | 0.002 * |
| Age of Initiation | Smokescreen | Nicotinic | 0.078 | 0.028 | 0.003 * |
| Cigarettes per Day | Smokescreen | Nicotinic | 0.079 | 0.031 | 0.005 * |
| Smoking Initiation | Smokescreen | Nicotinic | 0.076 | 0.036 | 0.017 |
| Smoking Cessation | Smokescreen | Nicotinic | 0.061 | 0.028 | 0.015 |

Next, we used LDSC to estimate the total SNP-heritability (*h*^2^_*SNP*_) for each smoking phenotype. Total *h*^2^_*SNP*_ (SE) (on liability scale for the binary phenotypes, cessation and initiation) for each trait was smoking cessation: 0.074 (0.005); smoking initiation: 0.095 (0.003); age of initiation: 0.041 (0.002); cigarettes per day: 0.061 (0.002); and drinks per week: 0.049 (0.002). These estimates are similar to those estimated with the full (including 23andMe) dataset reported by Liu et al. (2019).

To assess the proportion of the estimated *h^2^_SNP_* attributable to each gene set, we applied partitioned LDSC. The heritable contribution of the Smokescreen gene set was significantly enriched for age of initiation, cigarettes per day, smoking cessation, and smoking initiation (enrichment, the proportion of *h*^2^_*SNP*_ attributable to the gene set divided by the proportion of SNPs within the gene set, ranged from 1.4-2.3; Table 2). The Nicotinic gene set was significantly enriched only for smoking cessation. Although Nicotinic gene set enrichment estimates for all phenotypes were comparatively high (ranging from 1.59-3.27; Table 2), the relatively small number of genes, and therefore small number of variants as a proportion of all variants, led to high uncertainty in these estimates. The Nicotinic gene set enrichment standard errors were generally the largest and limited our statistical power to detect enrichment (Table 2). None of the control gene sets were significantly enriched for any trait. Overall, the enrichment analyses indicated that the genes present on the Smokescreen array are indeed enriched for smoking-relevant genetic variants, and that at least some of the genetic signal in smoking phenotypes comes from variants within the nicotinic receptors and genes they interact with. However, these gene sets cumulatively explain a small percentage of *h*^2^_*SNP*_; less than 3% of *h*^2^_*SNP*_ is attributable to Nicotinic genes, and less than 13% is attributable to all of the Smokescreen genes (Table 2).

**Table 2.** LDSC heritability enrichment of each gene set annotation within the full baseline + LD model for the four smoking phenotypes and one control trait across the two smoking gene sets and three control gene sets. * indicates p < 0.01, the within-trait Bonferroni correction for multiple testing.

| Trait | Gene Set | Proportion of SNPs in annotation | $h^2_{\text{SNP}}$ proportion | $h^2_{\text{SNP}}$ proportion Std. Err. | Enrichment | Enrichment Std. Err. | p |
| --- | --- | --- | --- | --- | --- | --- | --- |
| Age of Initiation | Nicotinic | 0.008 | 0.016 | 0.004 | 1.840 | 0.467 | 0.075 |
| Cigarettes per Day | Nicotinic | 0.008 | 0.028 | 0.009 | 3.274 | 1.092 | 0.045 |
| Drinks per Week | Nicotinic | 0.008 | 0.013 | 0.002 | 1.593 | 0.240 | 0.014 |
| **Smoking Cessation** | **Nicotinic** | **0.008** | **0.022** | **0.005** | **2.640** | **0.628** | **0.010 \*** |
| Smoking Initiation | Nicotinic | 0.008 | 0.020 | 0.005 | 2.375 | 0.601 | 0.024 |
| **Age of Initiation** | **Smokescreen** | **0.054** | **0.098** | **0.013** | **1.811** | **0.242** | **0.001 \*** |
| **Cigarettes per Day** | **Smokescreen** | **0.054** | **0.121** | **0.019** | **2.238** | **0.356** | **0.001 \*** |
| Drinks per Week | Smokescreen | 0.054 | 0.079 | 0.013 | 1.471 | 0.236 | 0.049 |
| **Smoking Cessation** | **Smokescreen** | **0.054** | **0.075** | **0.006** | **1.401** | **0.114** | **<0.001 \*** |
| **Smoking Initiation** | **Smokescreen** | **0.054** | **0.100** | **0.016** | **1.851** | **0.295** | **0.006 \*** |
| Age of Initiation | Height | 0.030 | 0.027 | 0.006 | 0.914 | 0.212 | 0.686 |
| Cigarettes per Day | Height | 0.030 | 0.035 | 0.007 | 1.169 | 0.249 | 0.494 |
| Drinks per Week | Height | 0.030 | 0.029 | 0.004 | 0.973 | 0.137 | 0.846 |
| Smoking Cessation | Height | 0.030 | 0.030 | 0.004 | 1.020 | 0.135 | 0.884 |
| Smoking Initiation | Height | 0.030 | 0.027 | 0.008 | 0.912 | 0.255 | 0.731 |
| Age of Initiation | ALZ | 0.043 | 0.072 | 0.012 | 1.685 | 0.286 | 0.015 |
| Cigarettes per Day | ALZ | 0.043 | 0.039 | 0.007 | 0.910 | 0.160 | 0.573 |
| Drinks per Week | ALZ | 0.043 | 0.053 | 0.007 | 1.243 | 0.162 | 0.137 |
| Smoking Cessation | ALZ | 0.043 | 0.050 | 0.005 | 1.171 | 0.128 | 0.183 |
| Smoking Initiation | ALZ | 0.043 | 0.039 | 0.010 | 0.905 | 0.225 | 0.675 |
| Age of Initiation | IBD | 0.012 | 0.016 | 0.007 | 1.350 | 0.606 | 0.564 |
| Cigarettes per Day | IBD | 0.012 | 0.017 | 0.005 | 1.443 | 0.417 | 0.284 |
| Drinks per Week | IBD | 0.012 | 0.018 | 0.004 | 1.565 | 0.341 | 0.104 |
| Smoking Cessation | IBD | 0.012 | 0.020 | 0.007 | 1.713 | 0.575 | 0.217 |
| Smoking Initiation | IBD | 0.012 | 0.021 | 0.010 | 1.841 | 0.888 | 0.345 |

## DISCUSSION

We examined the genes on the Smokescreen Genotyping Array and those involved with nicotinic receptors to assess whether they are associated with smoking phenotypes, and the degree to which they contribute to variation in the behavior (*h*^2^_*SNP*_). Both sets of genes were significantly associated with at least some of the tested smoking phenotypes. This is in contrast to the previous study of nicotinic genes by Melroy-Greif et al.^13^. However, the GSCAN summary statistics utilized a sample an order of magnitude larger than was available earlier, suggesting that the previous lack of gene set associations was due to low power. Additional associations of the Nicotinic genes may be found with larger sample sizes, though the partitioned LDSC analyses suggest that the nicotinic receptors and the genes they interact with cumulatively account for only 1.6-2.8% of *h*^2^_*SNP*_. This implies in combination with other studies (e.g., Liu et al.^7^) that although the nicotinic receptors are influential in smoking behaviors and play a key physiological role in response to nicotine, much of the genetic variation in these phenotypes is due to other pathways, consistent with a highly polygenic model of these complex traits.

Smokescreen incorporated roughly ten times the number of genes compared to the Nicotinic receptors, and accordingly accounted for a larger proportion of *h*^2^_*SNP*_ up to 12% of the genetic variance tagged by genome-wide markers. However, much of the genetic variance in smoking phenotypes remains unaccounted for by these variants and genes, and must be attributable to genes and pathways not tagged by this custom array. Together, these results suggest that the larger number of smoking-relevant genes in the Smokescreen gene set as a whole provide limited additional information over nicotinic receptor-related genes. However, since there were numerous individual genome-wide significant genes uniquely belonging to either the Smokescreen or Nicotinic gene set (Suppl. Table S1), specific genes within each set are clearly still contributing to variation in smoking behaviors. For predictive purposes and individualized treatment for nicotine use, additional variants outside of classical ‘addiction genes,’ including nicotinic receptors, will need to be considered.

Our study was limited to the GSCAN sample^7^, restricted to only those of European ancestry, and no independent replication samples are available. Furthermore, while GSCAN contains the currently largest available sample for summary statistics, we were limited to those excluding 23andMe data, and increasing the sample size would likely increase our power to detect gene set associations. However, this is unlikely to substantially change our conclusions, particularly as LDSC-based *h*^2^_*SNP*_ estimates indicate that these genes collectively account for a small proportion of the genetic variance. Notably, the GSCAN performed GWAS of common variants only, and thus the gene set associations and *h*^2^_*SNP*_ estimates presented here are restricted to the influence of common variants. Rare variants within genes of these gene sets may influence smoking behaviors and therefore these genes may yet have a larger role in smoking behavior than these analyses suggest. However, even if this is the case, it is unlikely that rare variant *h*^2^_*SNP*_ would account for the large discrepancy between current total *h*^2^_*SNP*_ estimates and twin-based *h*^2^.

Smoking is one of the leading cause of premature death in the United States^1^, and new forms of nicotine use, such as e-cigarettes, have gained prominence^3^. There remains, therefore, a critical need to understand the underlying biology of nicotine use, and identifying key genes and sets of genes is one possible avenue toward this goal. Nicotinic acetylcholine receptors play a key role as the binding sites for nicotine and the genes with which they interact influence smoking behaviors, as do genes traditionally thought of as ‘addiction’ genes. However, while these loci influence genetic variation in smoking behaviors, it is clear that these specifically targeted genes do not yet account for a large proportion of the heritability in smoking, and will therefore be of limited use for predictive purposes^20^. Ever-larger association studies, beyond GSCAN^7^, will be instrumental for this purpose, by examining genome-wide, unascertained markers in combination with improved statistical power as well as incorporating other ‘omics datasets and advanced methodologies to go beyond positional mapping, such as imputing genetically-regulated gene expression^21^ or integrating information from animal models of addiction.

## ACKNOWLEDGEMENTS

Funding came from The Institute for Behavioral Genetics to LME and MAE, NIAAA F32 AA027435 to ECJ, and NIDA T32 DA017637 support to WEM. We thank Mengzhen Liu and Scott Vrieze for providing GSCAN summary statistics.

